# Understanding cephalopod regeneration: impact of sexual maturation, injury severity and general anesthesia on octopus arm restoration

**DOI:** 10.64898/2026.08.29.747974

**Authors:** Samraggi Chakraborty, Łukasz Bijoch, Fan-Che Kung, Ping-Jui Hsieh, Zhen-Hong Yang, Kuo-Sheng Lee

**Author notes:** Corresponding author: Kuo-Sheng Lee.

## Abstract

Octopuses exhibit remarkable regenerative capacity, as shown by their ability to fully restore a whole arm. During this process, a complex appendage containing muscle, connective tissue, vasculature, suckers, sensory structures, peripheral nerves and the axial nerve cord (ANC) is regrown. Although the morphology of octopus regeneration is increasingly well described, the physiological variables that determine regenerative speed, including the potential contribution of ion-channel-dependent processes, remain poorly understood. In this work we examined the influence of reproductive state, injury severity, biological sex and magnesium chloride (MgCl_2_) anesthesia, on the rate of longitudinal arm regrowth. Reproductive stage was the strongest determinant of regenerative performance: pre-reproductive stage (pre-RS) animals showed robust regrowth, whereas reproductive stage (RS) animals regenerated significantly slower. In pre-RS animals, regeneration speed scaled strongly with injury severity, whereas this relationship was absent in RS animals. Biological sex did not independently predict regeneration speed, and MgCl_2_ exposure showed no long-term inhibitory effect on regeneration. Our final model explained 81.4% of variance in regeneration speed, with reproductive state and injury severity having major effects. These findings demonstrate that sexual maturation and optic-gland-mediated senescence act as an irreversible physiological block against somatic regeneration, and provide empirical support for mechanosensory and bioelectric-dependent scaling models of cephalopod limb restoration.

## Introduction

Octopuses possess an exceptional capacity to completely regenerate functional arms following traumatic injury, autotomy, or surgical amputation (Kier, 2016; Imperadore & Fiorito, 2018). Unlike vertebrate limbs that are supported by rigid articulated endoskeletons, the cephalopod arm functions as a continuous muscular hydrostat, in which three-dimensional skeletal support and omnidirectional bending are generated through densely packed arrays of transverse, longitudinal, and circular/helical muscle fibers maintaining a constant tissue volume (Kier, 2016; Grasso, 2014). Functional limb replacement requires not only reconstructing complex soft tissues, including vasculature, connective tissue septa, and chromatophore-bearing dermal layers, but also regenerating an intricate distributed nervous system. This apparatus comprises thousands of serial suckers equipped with chemotactile and mechanosensory receptors, four intramuscular nerve cords (INCs), and a continuous ganglionated axial nerve cord (ANC) that coordinates local sensorimotor reflex loops (Gutfreund et al., 2007; Hochner, 2012; Olson & Ragsdale, 2023; Benedict et al., 2026). Functional recovery thus demands the coordinated re-establishment of excitable cellular membranes, precise sensory transduction pathways, axonal pathfinding, and peripheral motor control (Imperadore et al., 2019; Benedict et al., 2026).

Histological and ultrastructural investigations across the *Octopus vulgaris* species complex demonstrate that arm repair proceeds through a stereotypic sequence (Shaw et al., 2016; Imperadore et al., 2022). Immediately after amputation, wound healing initiates with rapid sphincter-like contraction of circular muscle bundles and near complete re-epithelialization across the amputated stump within 24–48 hours, shielding internal tissues without scar formation (Shaw et al., 2016; Antunes et al., 2013). This initial phase is succeeded by blastema emergence (weeks 1–2 post-injury), characterized by the accumulation of undifferentiated mesenchymal progenitor cells beneath the wound epithelium, followed by a period of intense cellular proliferation and longitudinal elongation (weeks 3–4), and culminating in progressive morphogenesis, sucker organogenesis, and neural differentiation (>4 weeks) (Fossati et al., 2013; Imperadore et al., 2019, 2022). During this proliferative burst, classic signaling enzymes undergo dynamic spatiotemporal regulation. In particular, acetylcholinesterase (AChE) activity changes dynamically during regeneration, remaining low during the early proliferative/blastema phase, increasing during subsequent morphogenesis, and declining again as regeneration progresses (Fossati et al., 2013)

Beyond local biochemical cascades, somatic repair in cephalopods is modulated by sexual maturation and reproductive senescence (Wodinsky, 1977; Anderson et al., 2002; Di Cristo, 2013). This developmental program divides the octopus lifespan into two distinct physiological states: a pre-reproductive stage (pre-RS) focused on active predation, rapid somatic growth, and robust tissue regeneration; and a terminal reproductive/senescent stage (RS) characterized by metabolic catabolism, chromatophore pallor, behavioral lethargy, unhealed skin ulcers, and the apparent cessation of wound repair (Anderson et al., 2002; Holst, 2022; Roura et al., 2024). This maturation is governed by the optic-gland neuroendocrine axis, an endocrine signaling center functionally analogous to the vertebrate hypothalamic-pituitary axis (Wodinsky, 1977; Di Cristo, 2013; Wang et al., 2022). Historical experiments demonstrated that surgical removal of the optic glands in mated or brooding octopuses prevents post-reproductive mortality and resumes feeding, whereas experimental activation of the optic glands accelerates gonadal maturation at the expense of somatic tissue maintenance and regeneration (Wodinsky, 1977; O’Dor & Wells, 1978; Imperadore & Fiorito, 2018). Recent transcriptomic and lipidomic analyses have unraveled the molecular basis of this systemic switch: sexual maturation triggers massive optic gland upregulation of steroidogenic and cholesterol pathways, producing surges of progesterone, 7-dehydrocholesterol, and bile acid precursors that drive widespread somatic decline, muscle proteolysis, severe anorexia, and eventual maternal death (Wang & Ragsdale, 2018; Wang et al., 2022).

Prior work has established the basic stages of regeneration and identified reproductive maturation and senescence as major suppressors of somatic repair, but the extent to which regeneration scales with the magnitude of injury, and whether this scaling is preserved across life-history states, has not been systematically tested. In this work, we perform longitudinal measurements of octopus arm regeneration and we test how reproductive state, biological sex, injury severity, and MgCl_2_ anesthesia shape regenerative dynamics.

## Materials and Methods

### Animals and husbandry

All experimental procedures and animal housing protocols were reviewed and formally approved by the Institutional Animal Care and Use Committee (IACUC) of the Institute of Biomedical Sciences, Academia Sinica, Taiwan (Protocol Approval Number: 26-03-3081). All procedures conformed to international consensus guidelines for the accommodation, care, and welfare of cephalopods in research (Andrews et al., 2013; Fiorito et al., 2015).

Wild-caught East Asian common octopuses *(Octopus sinensis d’Orbigny, 1841; member of the Octopus vulgaris species complex; Gleadall, 2016)* were housed individually in a dedicated marine biosecurity facility operating a closed-loop recirculating aquaculture system. Routine target conditions were 21–25 °C, pH 8.0–8.5, salinity approximately 33‰, ammonia and nitrite near zero, and nitrate below 40 mg L^−1^. Animals were daily offered marine prey (grass shrimps, clams, fishes) according to routine husbandry practice, consistent with previously reported octopus husbandry protocols (Andrews et al., 2013; Benedict et al., 2026).

The reproductive stage of each octopus was tracked longitudinally based on standardized physiological, morphological, and ethological criteria (Anderson et al., 2002; Di Cristo, 2013; Wang et al., 2022). Pre-RS individuals were characterized by active predatory hunting, rapid prey capture, dynamic chromatophore patterning, strong sucker adhesion, and regular exploratory locomotion. In contrast, RS individuals showed severe anorexia (food refusal), loss of muscle tone, permanent skin pallor, and non-healing epidermal lesions. RS are also characterized by progressive apathy, erratic wandering, severe cachexia (Anderson et al., 2002; Wang et al., 2022; Roura et al., 2024). Individuals transitioning from pre-RS to RS during the experiment were tracked and their exact transition weeks were recorded.

### Arm injury and longitudinal morphometry

Distal arm amputations (1 to 3 arms per animal; N = 33 arms across 23 individuals) were performed at calibrated injury severities corresponding to 10%, 30%, 50%, or 70% of total arm length (measured from the oral-web margin to the distal arm tip) across anterior, posterior, left, and right arm pairs (L1–L4, R1–R4) (Shaw et al., 2016; Imperadore et al., 2019). For animals receiving general anesthesia, immersion in specialized anesthetic seawater containing 110–120 mM MgCl2 was conducted in a dedicated aerated chamber (Messenger et al., 1985; Brophy et al., 2018; Butler-Struben et al., 2021). Octopuses were monitored continuously until achieving deep surgical anesthesia (characterized by slowed ventilation <5 breaths/min, bilateral pupil dilation, flaccid muscular relaxation, and complete loss of arm withdrawal reflexes upon tactile forceps pinch) (Messenger et al., 1985; Brophy et al., 2018; Butler-Struben et al., 2021; Fiorito et al., 2015). Surgical amputations were conducted under sterile conditions with continuous oxygenated seawater mantle irrigation using precision micro-scalpel and surgical scissors (Shaw et al., 2016). Following surgery, animals were revived in pure oxygenated seawater before being returned to their home aquaria.

Longitudinal arm regrowth was measured weekly across an 84-day post-operative observation. At each weekly interval, the regenerating arm was gently positioned flat against a submerged acrylic metric grid board (1 mm grid divisions) and photographed. Longitudinal regrowth length was quantified from the initial amputation plane to the distal-most tip of the blastema using ImageJ/Fiji. Daily regenerative elongation rate (mm/day) was calculated as Δlength / Δtime (Fossati et al., 2013; Imperadore et al., 2022).

### Statistical analysis

Bivariate group comparisons were performed using non-parametric Mann-Whitney U tests for two-group medians and Welch’s t-tests for unequal variances. Kruskal-Walls test was used to compare four groups. Bivariate scaling relationships between regenerative velocity and injury severity (%) were evaluated using Pearson’s product-moment correlation coefficient (r) and linear regression fits. Statistical significance was defined at two-tailed α = 0.05, and statistical significance was defined as p < 0.05.

To evaluate the independent contributions of biological and procedural predictors to arm regeneration, we formulated a multivariable Ordinary Least Squares (OLS) linear regression model (Fox & Weisberg, 2019; Zar, 2010):

*Regeneration Speed (mm/day) = β0 + β1*·*(reproductive stage) + β2*·*(anesthesia) + β3*·*(arm side) + β4*·*(sex) + β5*·*(amputation severity) + ε, where*

*β0* = *y*-intercept (baseline regeneration speed when all predictors equal 0)

*β1-5* = regression coefficients for each predictor variable

*ε = error term*

Categorical predictors were dummy-coded with reference baselines defined as: pre-RS (0 = pre-RS, 1 = RS), (-)MgCl_2_ (0 = (-)MgCl_2_, 1 = (+)MgCl_2_), female (0 = female, 1 = male), and left arm side (0 = left, 1 = right). Amputation severity was entered as a continuous covariate (10% to 70%). Model goodness-of-fit was evaluated by total R^2^, Adjusted R^2^, and omnibus F-statistic.

## Results

### Arm longitudinal regeneration is heterogeneous and reproductive stage dependent

Octopuses were classified as pre-RS and RS based on previously described behavioral and physical characteristics (see Methods; Anderson et al., 2002). To characterize the regeneration time course of octopuses at different stages, we dissected arm tissue from octopuses of both sexes and we tracked longitudinal arm regrowth weekly over an 84-day post-amputation period (Fig. 1A-C).

**Figure 1:**
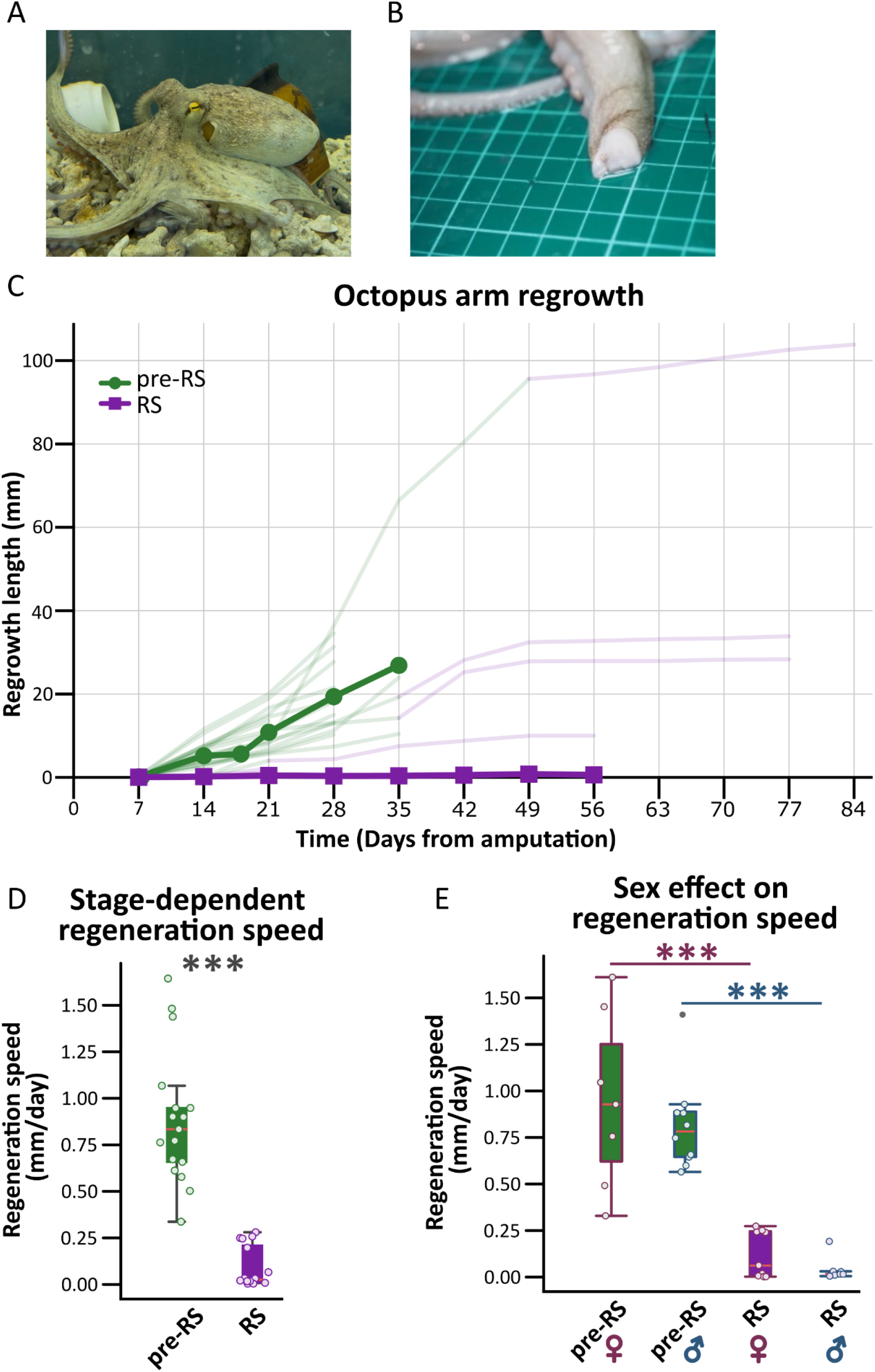
Arm regeneration of octopuses at different reproductive stages. **A)** Photography of *Octopus sinensis*. **B)** Photography of a dissection site on a grid table for measurement. **C)** Graph showing arm regrowth of pre-RS (green) and RS (purple) octopuses. Thick lines show group means and thin lines individual arms. Some animals changed their reproductive stage during the experiment, which is marked by changes in thin lines colors (green to purple). After the transition, arms were excluded from averaging. **D)** Box plots showing the average regeneration speed for pre-RS (green) and RS (purple) groups. **E)** Box plots showing the average regeneration speed for pre-RS females and males and RS females and males groups. Green and purple colors correspond to pre-RS and RS. Plump and blue colors correspond to females and males.

All pre-RS octopuses exhibited continuous, and progressive longitudinal arm elongation (Fig. 1C). Individual arm trajectories displayed an initial latency interval of 7 days with minimal measurable elongation but with observable wound contraction and blastema knob emergence. This was followed by sustained, accelerated linear growth from Day 14 through Day 84 (reaching up to >100 mm total regrowth; Fig. 1C). Across the pre-RS cohort (N = 17 arms), the mean daily regeneration velocity was 0.886 ± 0.089 mm/day (median 0.834 mm/day, IQR: 0.612–1.067 mm/day; Fig.1D). On the contrary, RS octopuses exhibited profound regenerative suppression, with arms failing to mount sustained outgrowth despite achieving initial wound epithelialization (mean 0.091 ± 0.024 mm/day, median 0.021 mm/day, IQR: 0.008–0.247 mm/day; Fig. 1C-D).

Three pre-RS octopuses with four operated arms underwent sexual maturation during the experiment and transitioned into the RS stage (Fig. 1C; indicated by thin line color shifts from green to purple). Overall, the transition did not immediately slow arm regeneration, and a plateau emerged only several weeks after the transition. This delayed slowdown likely reflects a prolonged transitional phase. Post-transition measurements were excluded from pre-RS averages.

We next stratified regeneration velocities by biological sex across both reproductive stages (Fig. 1E). In the pre-RS group, the raw difference of −0.104 mm/day between sexes was minor and not statistically significant (Fig. 1E).

Number of groups (arms): pre-RS females (N = 7), pre-RS males (N = 10), RS females (N = 9), and RS males (N = 7). Statistical difference for **D** was calculated by Mann-Whitney test, with p < 0.0001. And for E with Kruskal-Wallis test (post-hoc: Mann–Whitney and Bonferroni tests) with p < 0.0001 and pre-RS female vs. RS-female p = 0.00105 and pre-RS male vs. RS male p = 0.00062. For **B** and **C**, the red lines represent the median of each group. The lower and upper borders of the box mark the 25th and 75th percentiles defining the interquartile range (IQR). The whiskers extend to 1.5 × IQR, and individual arm data points are overlaid as dots. *** = p < 0.0001

### Regeneration rate scale with injury severity in pre-RS octopuses

Injury severity provides an additional test of regenerative regulation, as higher loss requires more resources and scales with arm complexity. Yet, arm regrowth must proceed quickly enough to be functionally rebuilt during octopus’s short lifespan. Therefore, we tested whether regenerative growth is actively matched to the extent of tissue loss by correlating arm regeneration with procedure severity (10% to 70% of arm loss) across both reproductive stages (Fig. 2). In pre-RS octopuses, regeneration rate displayed a strong, statistically significant positive linear correlation with amputation severity (r = 0.73, p < 0.001). Pre-RS animals subjected to a minor 10% distal amputation regenerated at an average rate of 0.56 ± 0.06 mm/day, whereas animals suffering a massive 70% amputation accelerated their regeneration velocity nearly three-fold to 1.52 ± 0.11 mm/day.

**Figure 2:**
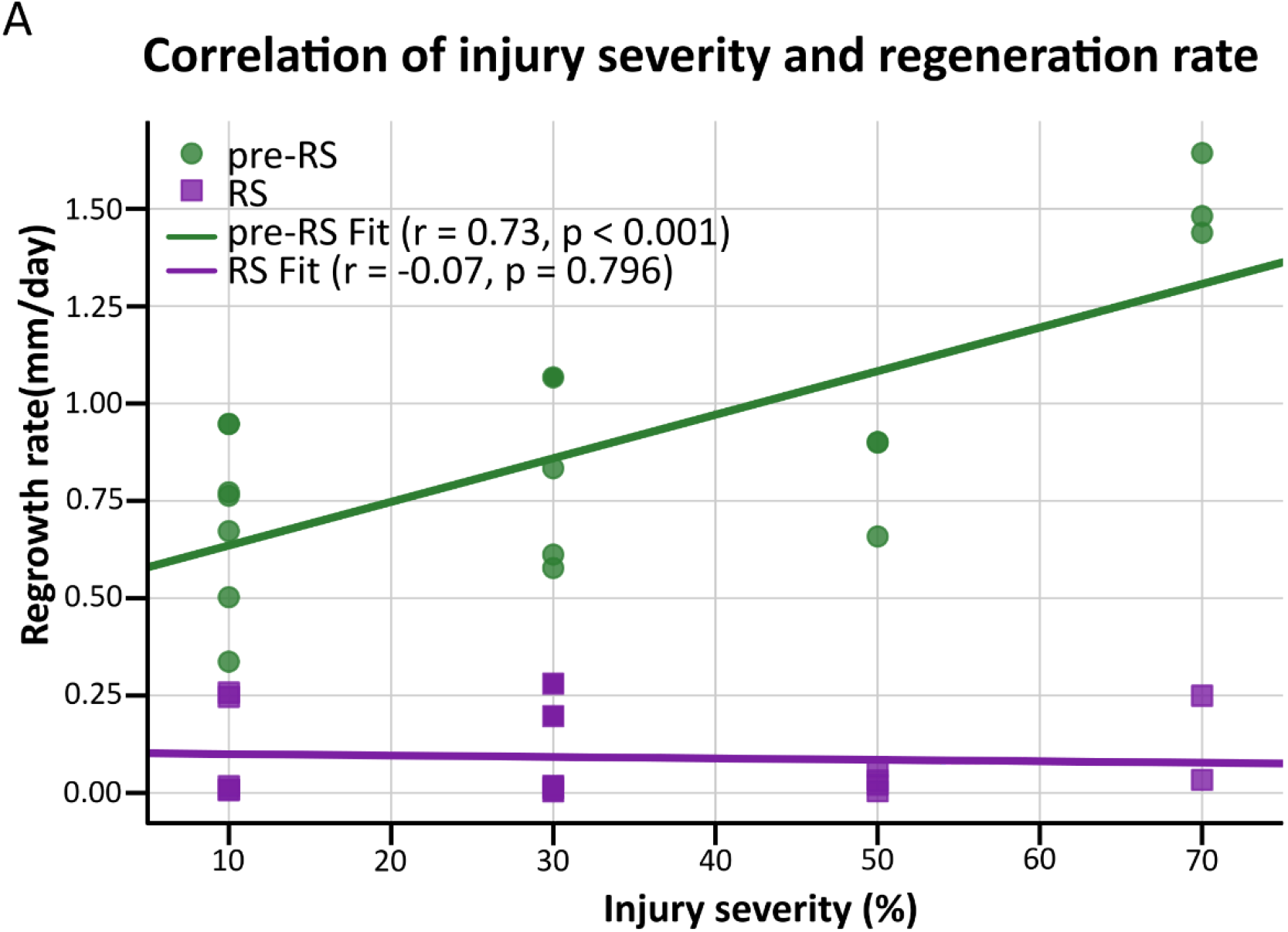
Correlation of injury severity and regeneration rate. **A)** Pearson’s correlations of injury severity and regeneration rate in pre-RS (green) and RS (purple) animals.

Contrary, RS octopuses remained entirely confined to a low regenerative regime across all injury magnitudes, exhibiting no correlation between tissue loss severity and regrowth velocity (r = -0.07, p = 0.796). Therefore we conclude that compensatory injury-severity acceleration is an active physiological adaptation that operates robustly in healthy pre-reproductive octopuses but is completely extinguished upon entry into reproductive senescence.

### Peri-operative MgCl2 general anesthesia does not inhibit long-term regeneration

MgCl_2_ is widely used as a cephalopod anesthetic that reversibly suppresses afferent and efferent neural signaling through changes in ionic concentration (Messenger et al., 1985; Brophy et al., 2018). We tested whether peri-injury exposure, occurring while the wound was still exposed, influenced subsequent regeneration (Fig. 3). We therefore stratified our data by reproductive stage (pre-RS vs. RS) and by anesthesia administered (Respectively: (+)MgCl_2_ and (-)MgCl_2_ ; Fig. 3A).

**Figure 3:**
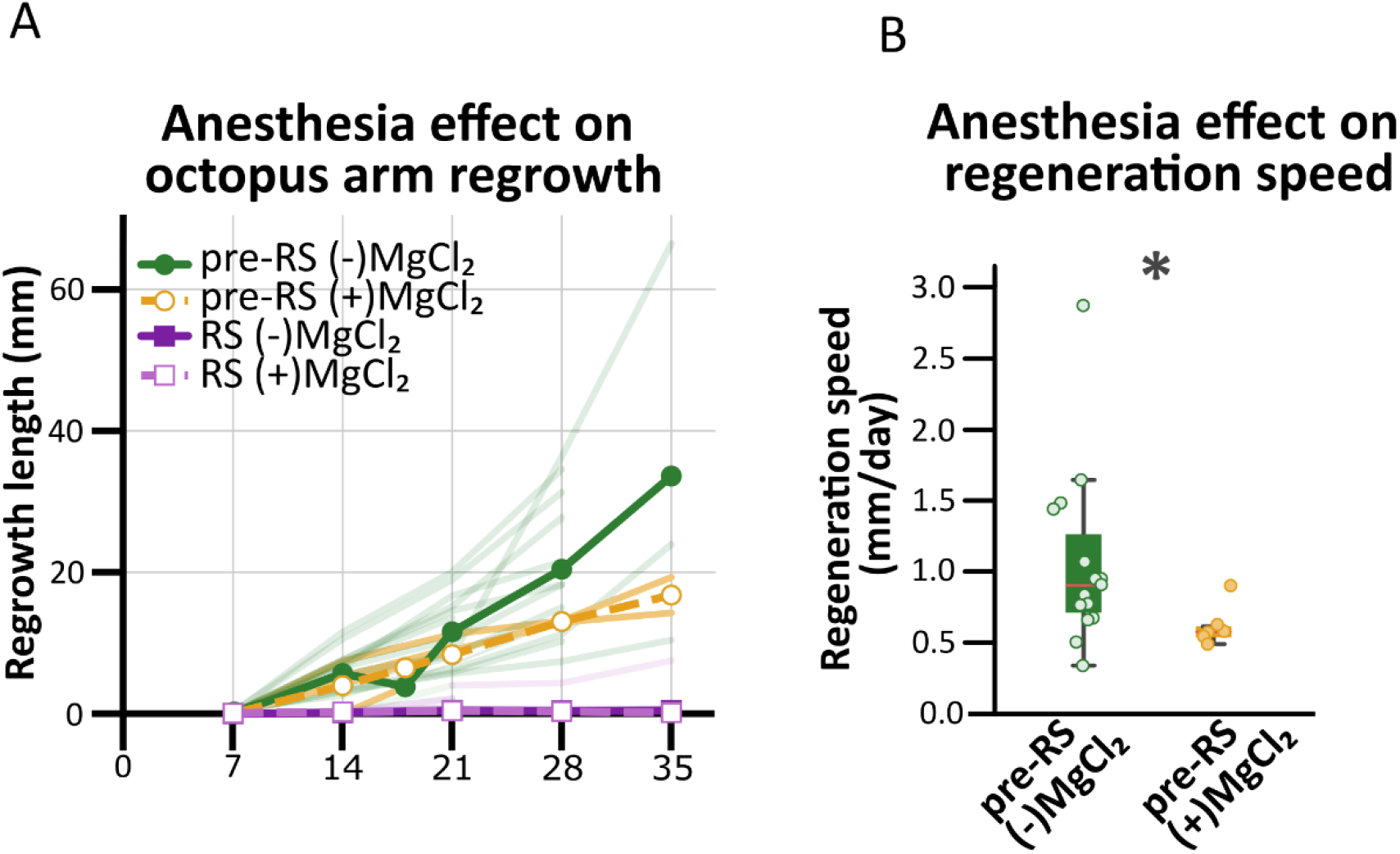
Regeneration of arms dissected with or without MgCl_2_ anesthesia. **A)** Graph showing arm regrowth of pre-RS and RS octopuses in different anesthesia conditions. Thick lines show group means and thin lines individual arms. Colors of groups are presented on the top left corner of the graph. Thick lines represent cohort means; faint lines show individual trajectories. **B)** Box plots showing the average regeneration speed for pre-RS with MgCl_2_ anaesthesia (green) and without MgCl_2_ anaesthesia (orange).

We found that anesthesia slightly affects regrowth rate of pre-RS animals, with no effects on RS octopuses (Fig. 3A). In raw bivariate comparisons within the pre-RS cohort, (-)MgCl_2_ arms exhibited a higher mean regrowth rate (0.997 ± 0.115 mm/day) than arms exposed to MgCl_2_ anesthesia (0.609 ± 0.061 mm/day; Fig. 3B). However, this effect could be cofounded by variety of amputation severity: arms in the pre-RS (+)MgCl_2_ group were subjected to smaller amputations (mean severity 18.3 ± 3.1%) compared to (-)MgCl_2_ (mean severity 42.0 ± 5.1%).

Number of groups: N MgCl_2_ (+) = 6 and N MgCl_2_ (-) = 15. For **B** the red lines represent the median of each group. The lower and upper borders of the box mark the 25th and 75th percentiles defining the interquartile range (IQR). The whiskers extend to 1.5 × IQR, and individual arm data points are overlaid as jittered black dots. Dots represent data for individual arms. Statistical difference for B was tested with Welch’s t-test, p = 0.019.

### The multivariable model explains most observed variation

To integrate all biological, anatomical, and experimental predictors into a unified predictive framework, we estimated a multivariable OLS regression model (R^2^ = 0.814, Adjusted R^2^ = 0.780, F(5, 27) = 23.63, p = 1.08 × 10^−9^; Fig. 4 & Table 1). The model identified reproductive stage as the dominant negative predictor of regeneration velocity (β = -0.807, SE = 0.089, t = -9.080, p = 1.08 × 10^−9^, 95% CI: [-0.989, -0.624]) and amputation severity as a highly significant positive driver of regrowth (β = +0.00624 per % loss, SE = 0.00191, t = 3.274, p = 0.0029, 95% CI: [0.00233, 0.01016]).

**Table 1:** Multivariable OLS regression model for arm regeneration velocity. Model R^2^ = 0.814, Adjusted R^2^ = 0.780, F(5, 27) = 23.63, p = 1.08 × 10^−9^. N = 33 arms across 23 individuals. Continuous predictor: Amputation Severity (10% to 70%). Baseline reference categories: Pre-RS, Female, Unanesthetized Control, Left Arm Side.

| Variable | Coefficient (Beta) | Standard Error | p-value | 95% CI Lower | 95% CI Upper |
| --- | --- | --- | --- | --- | --- |
| Intercept (baseline: pre-RS, (-)MgCl<sub>2</sub>, female, left) | 0.7546 | 0.1037 | 7.89e-08 | 0.5419 | 0.9674 |
| Amputation severity (%) | 0.00624 | 0.00191 | 0.0029 | 0.00233 | 0.01016 |
| Arm side (right) | 0.0055 | 0.0855 | 0.9488 | -0.1699 | 0.1810 |
| Anesthesia (MgCl<sub>2</sub>) | -0.0437 | 0.0963 | 0.6539 | -0.2413 | 0.1540 |
| Biological sex (male) | -0.1127 | 0.0835 | 0.1886 | -0.2841 | 0.0587 |
| Reproductive Stage (RS) | -0.8066 | 0.0888 | 1.08e-09 | -0.9888 | -0.6243 |

**Figure 4:**
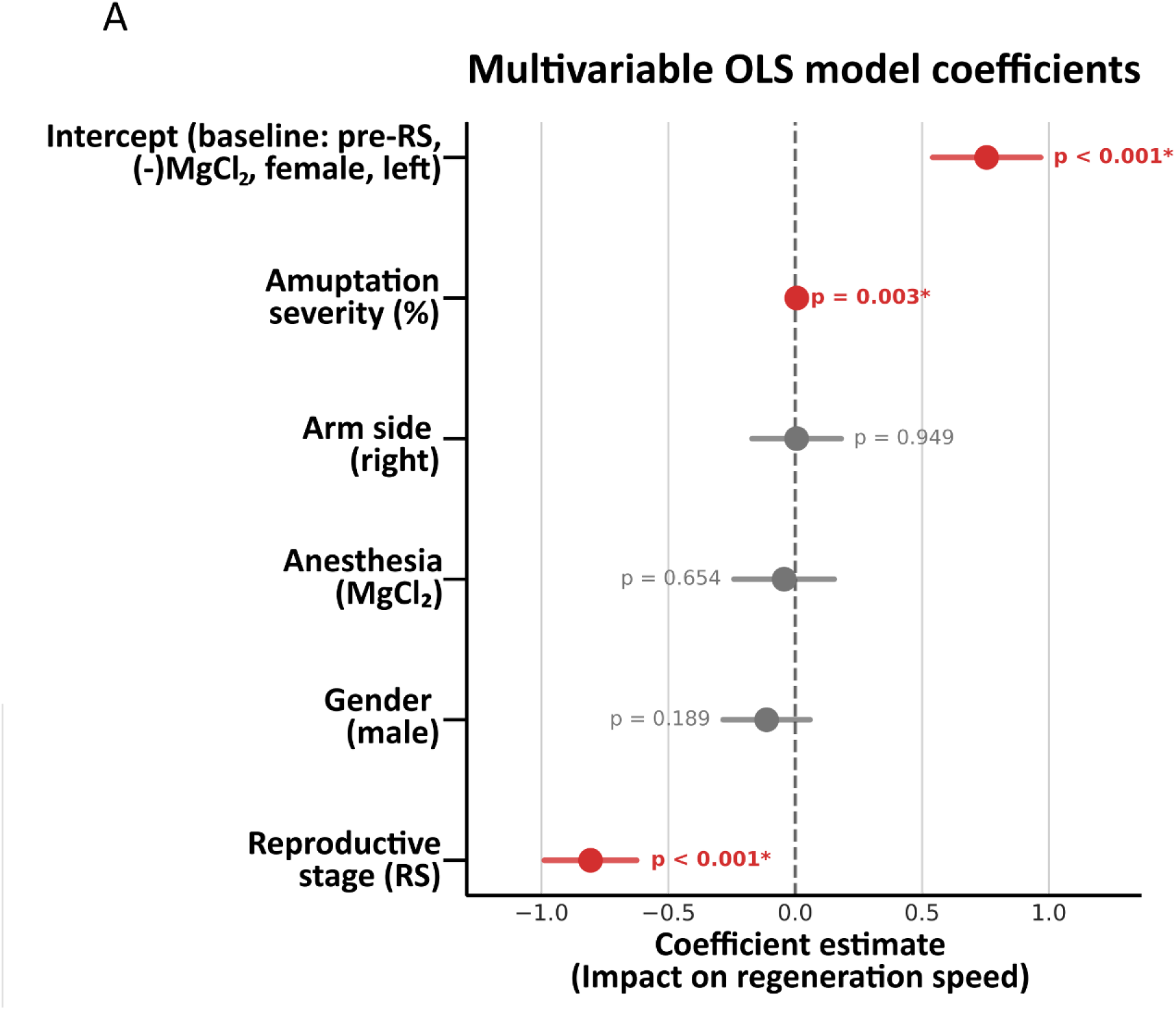
Multivariable OLS Model and predictor coefficients. Forest plot of standardized multivariable OLS regression coefficients (Beta ± 95% Confidence Intervals) predicting daily arm regeneration speed (mm/day; Total Model R^2^ = 0.814, F = 23.63, p < 0.001). Intercept (Baseline: Pre-RS, Female, Unanesthetized Control, Left arm): β = +0.755 (***p < 0.001); Reproductive Stage (RS): β = -0.807 (***p < 0.001); Amputation Severity (%): β = +0.00624 (**p = 0.0029); Biological Sex (Male): β = -0.113 (p = 0.189, NS); Anesthesia (MgCl2): β = -0.044 (p = 0.654, NS); Arm Side (Right): β = +0.0055 (p = 0.949, NS).

When evaluated in the multivariable OLS regression model controlling for covariances, MgCl2 anesthesia showed no significant independent effect on regeneration speed (β = -0.044, SE = 0.096, t = -0.453, p = 0.654, 95% CI: [-0.241, 0.154]). Furthermore, neither the biological sex (Male vs. Female: β = -0.113, p = 0.189), nor anatomical arm side (Right vs. Left: β = +0.0055, p = 0.949) had an effect on octopus regeneration. Collectively, these results confirm that reproductive maturation state and injury magnitude are the primary determinants governing octopus arm regeneration.

## DISCUSSION

In natural marine environments, cephalopod arm damage and autotomy are extraordinarily prevalent, with ecological surveys indicating that approximately 50% of wild octopuses within the *Octopus vulgaris* species complex exhibit missing, damaged, or regenerating arms (Imperadore & Fiorito, 2018; Roura et al., 2024). While foundational histological staging was established in Mediterranean *Octopus vulgaris* (Shaw et al., 2016; Imperadore et al., 2019, 2022), our findings provide the first systematic longitudinal quantification of arm regenerative scaling and reproductive gating in the East Asian common octopus (*Octopus sinensis*; Gleadall, 2016). During the juvenile and subadult pre-reproductive period, somatic arm restoration is an indispensable physiological adaptation for predatory foraging, tactile manipulation, benthic camouflage, and predator defense (Kier, 2016; Grasso, 2014; Hochner, 2012). Our longitudinal findings demonstrate that pre-RS octopuses regenerate arms robustly, averaging nearly 1 mm of longitudinal outgrowth per day (Fig. 1). However, upon entry into sexual maturity and the RS, this potent regenerative capacity is dampened and eventually completely extinguished (Fig. 1B).

The transition into reproductive senescence is strongly regulated by the optic-gland neuroendocrine axis, which orchestrates systemic changes in feeding, metabolism and somatic maintenance (Wodinsky, 1977; O’Dor & Wells, 1978; Di Cristo, 2013). In semelparous cephalopods, sexual maturation triggers an irreversible endocrine shift away from somatic investment and tissue maintenance toward terminal reproductive output (Wodinsky, 1977; Wang & Ragsdale, 2018). Recent transcriptomic, lipidomic, and mass-spectrometric analyses have shown that activation of the optic gland upregulates extensive steroidogenic enzyme networks, producing elevated levels of progesterone, 7-dehydrocholesterol metabolites, and bile-acid intermediates that drive systemic muscle catabolism, severe anorexia, immune suppression, and somatic self-degradation (Wang & Ragsdale, 2018; Wang et al., 2022).

Despite comprehensive qualitative descriptions of cephalopod wound healing, a fundamental quantitative question remains: does arm regeneration velocity scale proportionally with the magnitude of tissue loss, and is this homeostatic scaling preserved across life-history stages (Imperadore & Fiorito, 2018)? In non-senescent octopuses with short lifespans (1–2 years), regenerating a severe injury (e.g., 70% arm loss) at the same baseline velocity as a minor injury (e.g., 10% loss) would result in permanent functional impairment before sexual maturity. Biomechanical and bioelectrical models predict that larger tissue loss alters stump geometry, increases mechanical tension across the wound, and generates sustained bioelectric injury potentials that could scale the mitotic recruitment of progenitor cells (Levin, 2007, 2021; Cervera et al., 2018). Here we discovered that arm regeneration rate is not fixed, but scales strongly and positively with the magnitude of tissue loss. Octopuses subjected to a severe 70% amputation accelerated their regenerative elongation nearly three-fold compared to those with a 10% injury (1.52 vs. 0.56 mm/day). In short-lived semelparous organisms (lifespan 12–18 months), this homeostatic acceleration represents a vital evolutionary adaptation, ensuring that even catastrophic arm loss can be functionally restored prior to the onset of the terminal reproductive window (Imperadore & Fiorito, 2018; Roura et al., 2024). A mechanosensitive pathway could contribute to the graded response observed in pre-RS animals. In cephalopods, specialized mechanosensitive ion channels (TRPN), nociceptive (TRPV) and mechanogated Piezo cation channels (van Giesen et al., 2020; Allard et al., 2023; Pieroni et al., 2026; Coste et al., 2010; Imperadore et al., 2022) represent plausible molecular transducers linking physical strain to cellular regenerative output. Calibrated stretch or indentation combined with calcium imaging or electrophysiology would allow this possibility to be tested without assigning causality to a specific family prematurely.

Magnesium chloride (MgCl2) is commonly utilized as a general anesthetic for cephalopod research (Messenger et al., 1985; Fiorito et al., 2015; Brophy et al., 2018). Mechanistically, elevated Mg2+ concentrations reversibly suppress neural signaling and muscular contraction by competitively inhibiting presynaptic voltage-gated calcium channels, thereby blocking neurotransmitter exocytosis at central and neuromuscular synapses and hyperpolarizing excitable membranes (Messenger et al., 1985; Butler-Struben et al., 2021). Because peri-operative anesthesia exposes stump tissues directly to hypermagnesemic seawater, concerns have existed regarding potential lingering toxicity on regenerating blastema cells. After adjustment for the major biological covariates, however, MgCl_2_ was not significantly associated with regeneration speed (β = -0.044, p = 0.654). After adjusting for baseline amputation severity, animals anesthetized with MgCl2 regenerated at rates indistinguishable from unanesthetized controls, confirming that standard MgCl2 protocols provide effective surgical analgesia and neuromuscular relaxation without compromising long-term cephalopod regenerative biology.

## Conclusions

Octopus arm regeneration is strongly regulated by reproductive state and only pre-RS animals regenerate. Pre-RS animals regenerate robustly and regeneration speed is correlated with injury severity. Biological sex and MgCl_2_ anesthesia do not show significant independent effects on regeneration rate.

## Limitations

Because multiple arm observations can originate from the same individual, the present arm-level analysis does not fully account for within-animal correlation. Future work should confirm these findings using verified animal identifiers within a clustered or mixed-effects modeling framework.

The final regression model explained 81.4% of observed variance, supporting the biological coherence of the current predictors while leaving room for body size, nutritional state, temperature, arm identity, wound geometry and regenerative phase.

## ACKNOWLEDGEMENTS

The authors thank Yu-Jung Su, Chun-Fang Lin and other laboratory members who contributed to octopus husbandry. The authors thank the Animal Core, the Pathology Core and the Common Facilities Core Laboratory at the Institute of Biomedical Sciences, Academia Sinica, Taiwan.

## AUTHOR CONTRIBUTIONS

S.C.: Data analysis, interpretation, writing. L.B.: Pilot imaging, supervision and writing. F.-C.K.: Conceptualization, methodology, arm measurements. P.-J.H. and Z.-H.Y.: Longitudinal photography and arm measurements. K.-S.L.: Conceptualization, supervision, writing and funding acquisition.

## DATA AVAILABILITY

All data needed to evaluate the conclusions in the paper are present in the paper and/or Supplementary Materials and will be shared by the lead contact upon request.

